# Off-Vertical Body Orientation Delays the Perceived Onset of Visual Motion

**DOI:** 10.1101/2022.11.13.516314

**Authors:** William Chung, Michael Barnett-Cowan

## Abstract

**Summary:** The integration of vestibular, visual and body cues is a fundamental process in the perception of self-motion and is commonly experienced in an upright posture. However, when the body is tilted in an off-vertical orientation these signals are no longer aligned relative to the influence of gravity. In this study, sensory conflict with the vestibular signal was introduced by manipulating the orientation of the body, generating a mismatch between body and vestibular cues due to gravity in the perception of upright and creating an ambiguous vestibular signal of either head tilt or translation. In a series of temporal-order judgment tasks, participants reported the perceived onset of a visual scene simulating rotation around the yaw axis presented in virtual reality with a paired auditory tone while in an upright, supine and side recumbent body positions. The results revealed that the perceived onset of visual motion was further delayed from zero (i.e., true simultaneity between visual onset and a reference auditory tone) by approximately an additional 30ms when viewed in a supine or side recumbent orientation compared to an upright posture. There were also no significant differences in the timing estimates of the visual motion between all the non-upright orientations. This indicates that the perceived timing of visual motion is negatively impacted by the presence of conflict in the vestibular and body signals due to the direction of gravity and body orientation, even when the mismatch is not in the direct plane of the axis of rotation.

## Introduction

Sensory conflict occurs when there is a discrepancy between multisensory signals. The mismatch of signals can arise due to differences in transduction and processing latencies between different sensory modalities. One such scenario where sensory conflict has been of interest is within the self-motion literature, where a mismatch between vestibular, visual and proprioceptive signals can commonly occur while using different vehicles of transportation and more recently with the application of virtual reality (VR). The vestibular system, responsible for detecting changes in acceleration of the head, is essential for providing us with information regarding our position and orientation in space and generates an internal sensation of self-motion (Angelaki and Cullen, 2008; Cheng and Gu, 2018). While vision provides similarly redundant information about self-motion through optic flow and vection (Palmisano et al., 2015), it additionally supplies us with information regarding the movement of objects in our environment. The integration of these cues usually creates a stable representation of self-motion, however when there is a mismatch between these signals there can be potential perceptual consequences such as disorientation, illusory self-motion, errors in decision making and in performing coordinated actions.

One of the primary reasons for the occurrence of conflict between visual and vestibular cues is that the vestibular signal is constantly providing feedback to the central nervous system. For example, when exposed to motion in VR while being physically stationary, visual feedback from the display signals self-motion while the vestibular system does not sense and signal any self-motion, leading to a mismatch. An additional source of conflict arising from the vestibular system is from the constant force of gravity on the hair-like structures found within the vestibular system known as stereocilia (Goldberg et al., 2012). The anatomy of the stereocilia enables the vestibular system to detect acceleration or changes in motion of the head, however the hair cells of the otolith organs of the vestibular system will detect not only translational acceleration but also the signal of acceleration related to the constant force of gravity. The influence of gravity can lead to ambiguous otolith signals from the vestibular system, where the acceleration feedback resulting from a head tilt can be the same as from a translational or rotational movement along the same axis [the tilt-translation ambiguity; (Crane, 2014)]. To differentiate the signals arising from these distinct events, the integration of otolith and semicircular canal signals (which detect rotational acceleration), along with visual and proprioceptive information is necessary to resolve the ambiguity of the vestibular signal.

A common method that has been used to explore the effect of gravity is through manipulating the orientation of the body. By changing the position of the body, the direction in which the gravitational vector affects the vestibular system is shifted accordingly since gravity is constant on Earth. Researchers have used this relationship between gravity and ambiguous vestibular signals in combination with visual motion to examine the influence of gravity on the perception of vection, the illusion of self-motion when the body is stationary. Much of the literature has focused on the difference in vection measures, such has magnitude and onset latencies, in an upright compared to supine, prone or recumbent position. When paired with different visual flow patterns, the gravitational vector can either support or conflict with the direction of vection depending on the pairing of sensory signals. Some findings in the literature have suggested that vection ratings improve when the gravitational vector is congruent with the direction of the visual motion in a supine posture (Kano, 1991; Young et al., 1986). However, there have also been reports showing that regardless of whether the direction of vection was with or against gravity, vection measures are higher in an upright posture (Guterman et al., 2012; Howard, 1987; Tovee, 1999; Watt, 1990).

Based on the current evidence, it appears to be more appropriate to describe the conflict between body orientation relative to gravity and visual motion not only in terms of the physical mismatch of the stimuli, but also considering the ecological aspect of being in a non-upright posture. The ecological view is a top-down approach based around what observers are accustomed to experiencing rather than focusing on the conflict that is physically present in the stimuli. In normal circumstances, people are typically exposed to motion when they are in an upright posture, thus the system is better tuned for perceiving specific types of motion along certain axes or planes. This ecological advantage of aligning the motion with its typical axis or plane has also been previously observed in studies for rotational vection (Howard, 1987; Tanahashi et al., 2012), vertical and horizontal vection (Guterman and Allison, 2019; Nakamura and Shimojo, 1998), vection onset latencies (Kano, 1991; Tovee, 1999) and heading discrimination (MacNeilage et al., 2010). It has also been shown that adding an oscillating effect to the visual display to simulate the bob and sway of natural locomotion increases vection strength regardless of body orientation and the resulting gravitational conflict (Guterman et al., 2012).

Another important ecological aspect for the integration of visual motion and body orientation is the congruency of the signals. As previously mentioned, the tilt-translation ambiguity of the vestibular system requires additional sensory feedback to be resolved and under common circumstances those additional sensory cues are typically consistent with the vestibular signal. For example, a forward translation is accompanied by an expanding optic flow, or alternatively a backwards head tilt is accompanied by an upwards tilt of the visual scene. However, in certain scenarios such as riding a vehicle or experiencing VR, the visual and vestibular signals can become incongruent, and researchers have been interested in exploring whether the change in this relationship between source of sensory information can affect our perception. In terms of body orientation, it has been reported that the perception of self-tilt is mainly processed in body coordinates (Crane, 2014), however the presence of visual scene tilt can improve or modulate that perception, but only when it is congruent with the tilt of the body (Di Cesare et al., 2014; Higashiyama and Koga, 2009). Furthermore, it has also been shown that body tilt can influence the perception of visual motion, including the perceived direction of an optic flow pattern (Bourrelly et al., 2010), the estimated distance travelled in VR (Kruijff et al., 2015) and the perceived realism of linear selfmotion (Groen and Bles, 2004). These findings indicate that there is a reciprocal relationship between visual and vestibular integration and highlight the importance of the congruence between the two signals.

To the best of our knowledge there has yet to be a study exploring the effects of body orientation on the perceived timing of motion. In the current experiment, we plan to examine the effects of sensory conflict on the perceived timing of visual motion during optic flow in the head-centered yaw rotation axis. The conflict will be introduced by altering the orientation of the head and body, such that the direction of visual motion will be either unaffected or incongruent with the relative vestibular signal due to gravity. In the upright posture, yaw rotations are invariant to the effects of gravity and the otolith signals which detect gravitational acceleration and is in the axis where yaw motion is commonly experienced. When in a supine or recumbent position, the otolith signals are consistent with signaling pitch or roll motions respectively and are not in the same rotational axis as the head-centered yaw visual motion presented to the observer. Our main hypothesis is that the perceived timing of visual motion should be less accurate and less precise when the visual flow is incongruent with the vestibular signal in supine and recumbent orientations compared to upright. We also explore whether there is a bias in the perceived timing for the direction of the visual motion in each of the body orientations. While the ambiguous tilt-translation signal is not in the axis of yaw rotations, the direction of gravity and visual motion are partially aligned in the lateral recumbent orientations (ex. rotations to the left are in the direction of gravity when in a left recumbent position). Thus, we hypothesize that there may be a difference between left and right rotations in the lateral recumbent positions compared to the upright and supine orientations due to the direction of gravity. Furthermore, we investigate whether there is an effect of a body tilt bias between the left compared to the right recumbent positions and also whether there are any differences in the off-vertical axes between lateral recumbent (left and right recumbent combined) compared to the supine orientation.

## Methods and methods

### Participants

Data was collected from twenty-four participants (11 males and 13 females; mean age = 26.9, SD = 3.7) recruited from the University of Waterloo. Five of the participants were volunteers, while the remaining participants were remunerated $10/h for their participation. All participants reported having no auditory or vestibular disorders and had normal or corrected-to-normal vision. This experiment was reviewed and received ethics clearance through a University of Waterloo Research Ethics Committee and all participants gave their informed written consent in accordance with the guidelines from the committee.

### Apparatus and Stimuli

The Oculus Quest system was used to provide the visual stimuli in this experiment. The Oculus Quest has a dual OLED display with a resolution of 1600×1440 (per eye) at 72 Hz refresh rate. The visual stimuli was a virtual starfield (Fig. 1b) randomly generated using the particle system built in Unity with the experimental control (trial randomization, recording of responses, graphical interface for monitoring progress of the experiment, etc.) programmed using the BiomotionLab Toolkit for Unity Experiments (bmlTUX) (Bebko and Troje, 2020). No visual fixation point or fixation instructions were provided to maintain a naturalistic experience and to prevent effects on self-motion perception (Garzorz and MacNeilage, 2017; Riecke and Schulte-Pelkum, 2013). Auditory stimuli were presented through in-ear headphones (Sennheiser CX 1.00).

**Figure 1.**
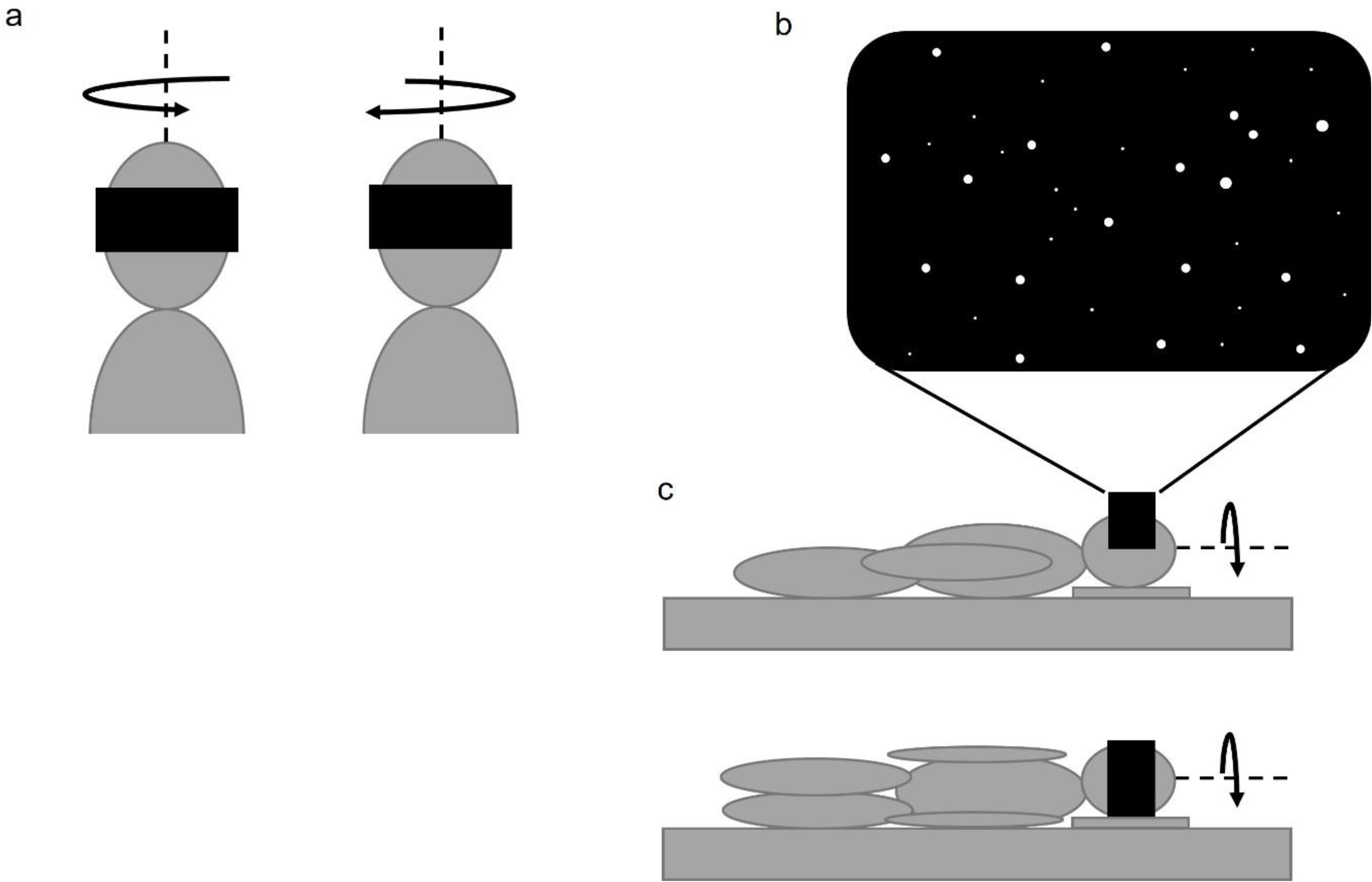
**a** Schematic depiction of an upright observer wearing an Oculus Quest VR HMD. Right and left rotations of the visual environment relative to the stationary observer were around the yaw axis indicated by the dotted line. **b** Starfield visual stimuli viewed using the VR HMD. **c** Schematic depiction of an observer wearing the VR HMD while either laying supine (above), or left side recumbent (below). Similar to the upright condition, right and left rotations of the visual environment relative to the stationary observer were around the yaw axis indicated by the dotted line.

In each trial, an 800 Hz tone was presented for 10 ms at nine stimulus onset asynchronies (SOAs) following a logarithmic scale: -600, -295, -145, -35, 0, 35, 145, 295, 600 ms, relative to the onset of the visual motion. Visual motion was a 10 degree angular rotation in the yaw axis in both left and right directions (Fig. 1a), with a peak velocity of 20 deg/s (Chung and Barnett-Cowan, 2022). A negative SOA indicates that the auditory stimuli was presented before the onset of visual motion, positive SOA indicates the auditory stimuli presented after the visual motion and zero indicates both stimuli occurring at the same time. The order of SOAs was randomized among trials and each SOA was presented nine times for both directions of visual motion for a total of 162 trials (9 SOAs x 9 repetitions x 2 rotation directions). Participants reported whether they perceived the onset of the visual motion occurring before or after the onset of the auditory tone using the buttons on a gamepad (Logitech Gamepad F310).

### Procedures

Participants were fitted with the Oculus Quest VR system before being positioned in one of four body orientations: upright, supine, left recumbent or right recumbent (Fig. 1c). All participants experienced every body orientation in a block design and the order of orientations were counterbalanced across participants. In the upright orientation, participants were seated on an office chair and in the remaining orientations an examination table was used with a foam pillow to keep participant’s head aligned with the body. The participants completed a temporal order judgment (TOJ) task in all four body positions, in which they had to indicate whether the onset of a visual motion stimuli (rotation of a starfield) or an auditory tone occurred first. The trials were self-paced in that the next trial would not begin until participants’ response with a button press on a gamepad was recorded. Participants were given a brief break (2 to 3 min) at the halfway point of each block and a longer break (5 min) in between blocks to minimize fatigue and maintain attention on the task.

### Data Analysis

Statistical analysis was conducted using SigmaPlot 12.5 and IBM SPSS Statistics 25. A logistic function which accounted for lapses in participants’ responses (Eq. 1) was fitted to the participants’ percentage of “visual motion first” responses as a function of the SOAs. Estimates of lapse rate, PSE and precision were derived from the three free-parameter equation to nine data points corresponding to the nine SOAs tested. The halfway (50%) inflection point of the sigmoid curve (x0) represents the point of subjective simultaneity (PSS) and the standard deviation of the function (b) as the just noticeable difference (JND) (Barnett-Cowan and Harris, 2011). A positive PSS score indicates that the motion had to occur before the auditory tone by the value of the score (in ms) to be perceived as occurring simultaneously, while a negative PSS score indicates that the motion had to occur after. In cases where the lapse function failed to appropriately fit the data (*R*2 < 0.2), an alternative 3-parameter logistic function (Eq. 2) was used instead. This substitution was resorted to 12 times out of a total of 192 fits and it yielded better R2 results that met the specified criteria in all the cases. We were unable to fit either function to the responses of three participants (*R*2 < 0.2 or *p* < 0.05) and were thus removed from the subsequent analysis.

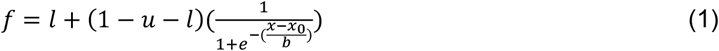

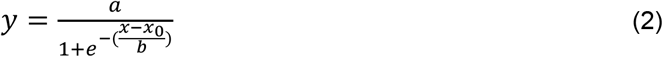

## Results

To observe whether there were any significant differences in the direction of the visual motion, paired t-tests were conducted between left and right rotations and revealed that there were no significant differences in the PSS between left and right rotations for all body orientations: upright [*t*(20) = 0.80, *p* = 0.44); supine [*t*(20) = 1.40, *p* = 0.18]; left recumbent [*t*(20) = 0.24, *p* = 0.81]; right recumbent [*t*(20) = -0.20, *p* = 0.84]. Thus, the data for each participant was collapsed across rotation direction and re-fitted with the logistic functions (Eqs 1 and 2) using the same methods as Xbefore. With the additional data points, we were able to fit the functions to all participants with seven substitutions using Equation 2 out of 96 total fits that yielded better *R*^2^ results meeting the specified criteria (*R*^2^ < 0.2). All subsequent data analyses were performed using the collapsed data, with two participants removed due to the fit of their responses failing to meet our criteria (*R*^2^ < 0.2 and *p* < 0.05) and an additional participant identified as a statistical outlier (over three times the interquartile range) (final sample n = 21; 9 males, 12 females; mean age = 27.5, SD = 3.3).

The average and standard error for the PSS and JND values are summarized in Table 1. One-sample t-tests were conducted to confirm that the PSS scores were significantly different from zero: Upright: [*t*(20) = 7.18, *p* < .001]; Supine [*t*(20) = 7.59, *p* < .001]; Left Recumbent [*t*(20) = 6.21, *p* < .001]; Right Recumbent [*t*(20) = 5.99, *p* < .001]. A paired t-test was conducted between the left and right recumbent orientations to determine whether there was an effect of the direction of the side lateral orientation on the participants’ response. The results showed that there were no significant differences between left and right recumbent orientations for both the PSS [*t*(20) = 0.29, *p* = 0.777, *d* = 0.063] and JND [*t*(20) = -0.18, *p* = 0.861, *d* = -0.039]. Since they were not significantly different, the values were then collapsed by taking the average of the individual PSS and JND values across the left and right recumbent orientations for each participant [PSS(M = 156.72, SE = 24.34); JND (M = 77.62, SE = 10.18)]. A subsequent Wilcoxon Signed-Rank Test between the collapsed lateral recumbent and supine orientation found no significant difference in PSS (*Z* = -0.82, *p* = 0.424, *r* = -0.203) and JND (*Z* = -1.20, *p* = 0.237, *r* = 0.299). Thus, the PSS and JND data for the lateral recumbent and supine orientations were consolidated into a single “Off-Vertical” orientation by averaging the lateral recumbent and supine data for each individual participant [PSS(M = 157.94, SE = 21.24); JND(M = 74.16, SE = 9.93)]. A final Wilcoxon Signed-Rank Test between the upright and off-vertical orientations revealed a significant difference in the PSS (*Z* = 2.42, *p* = 0.016, *r* = -0.602), indicating that off-vertically oriented participants required visual motion onset to precede the onset of an auditory reference tone significantly earlier (30ms earlier) than when they were upright. No significant difference for the JND was found (*Z* = -1.10, *p* = 0.281, *r* = 0.273) (Fig. 2).

**Table 1.** Summary of PSS and JND data across body orientations. Positive PSS values indicate that the visual motion needed to move prior to the auditory tone onset to be perceived as occurring simultaneously.

|  |  | Upright | Supine | Left Recumbent | Right Recumbent |
| --- | --- | --- | --- | --- | --- |
| PSS (ms) | Mean | 127.37 | 159.16 | 159.09 | 154.35 |
|  | SE | 17.74 | 20.98 | 25.64 | 25.78 |
| JND | Mean | 85.70 | 70.70 | 76.69 | 78.54 |
|  | SE | 15.44 | 13.78 | 10.94 | 11.93 |

**Figure 2.**
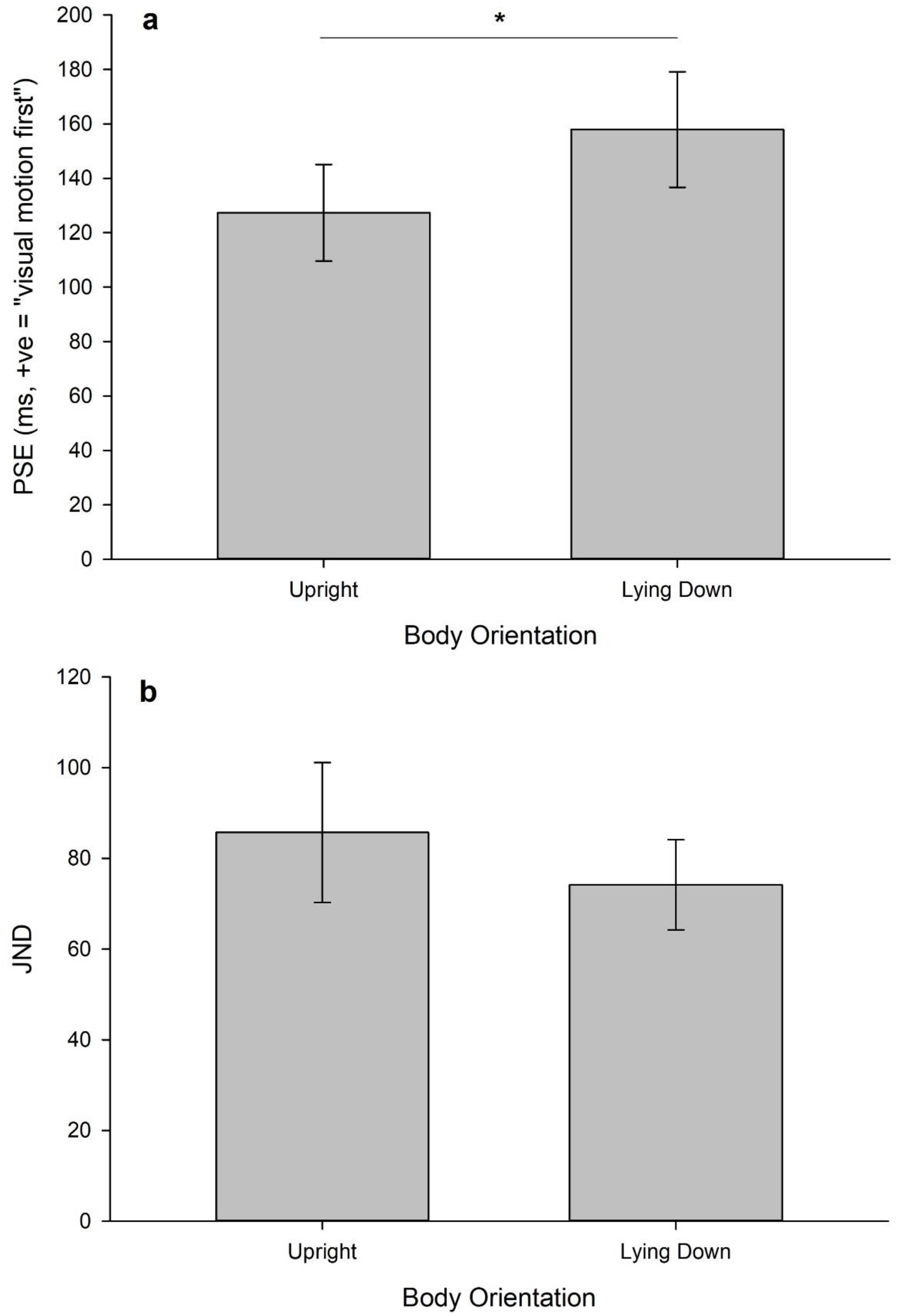
Summary of **a** point of subjective simultaneity (PSS), **b** just-noticeable difference (JND) between upright and lying down body orientation. Error bars are standard error.

## Dicussion

The purpose of this study was to explore the effects of varying body orientation on the perceived timing of visual motion. Changing the orientation of the body from upright dissociates the gravitational vector from the idiotropic vector or longitudinal body axis (Mittelstaedt, 1983) in signalling the direction of upright and we were interested in whether changes in this relationship would influence the perceived onset of visual rotation around the yaw axis presented in VR. The results showed that there were no significant differences in the direction of motion (left or right rotation), as well as between the various lying down orientations (left recumbent, right recumbent and supine). However, when all the lying down orientations were consolidated and compared to the upright orientation, the point of subjective simultaneity (PSS) was found to be significantly further from zero. The onset of the visual motion had to be presented an average of 158ms before the auditory tone for both stimuli to be perceived as occurring at the same time in the lying down orientations compared to only 127ms when upright. This suggests that there is a general attenuation effect on the ability to detect the onset of visual motion when the body is tilted relative to gravity resulting in sensory conflict between the vestibular and body signals. Contrary to previous findings that showed the precision in the perception of upright decreases when the body is tilted (Clemens et al., 2011; De Vrijer et al., 2009), there was no significant difference in the just-noticeable difference (JND) in the perceived onset of visual motion across all of the body orientations.

When in an upright posture, gravity and the longitudinal body axis are aligned and changes in the vestibular apparatus due to inertia signals the presence of self-motion. Motion is also commonly experienced in an upright posture, for example when walking or riding in a vehicle, and previous research has found that there is an ecological advantage for various aspects of visual motion in the direction of gravity and an upright orientation (Guterman and Allison, 2019; Guterman et al., 2012; Howard, 1987; Kano, 1991; MacNeilage et al., 2010; Nakamura and Shimojo, 1998; Tanahashi et al., 2012; Tovee, 1999, Watt, 1990). In our study, we found a significant difference in the perceived onset of visual motion when in an upright orientation compared to all the off-vertical orientations in which the timing estimate of visual motion was further then zero when the body was tilted in agreement with the ecological advantage hypothesis. Following this, we did not observe any differences between the supine and lateral recumbent orientations and a potential reason why may be due to the reference frame that the visual motion we presented was processed in. Our sensory systems encode information relative to different egocentric reference frames which need to be transformed into an allocentric representation that is gravity-centered to achieve a stable and upright representation of the world (Dakin and Rosenberg, 2018). For example, the visual system encodes the image on the retina in eye-centered coordinates, but when you change orientations your perception of verticality still remains intact to a certain a degree (i.e. a tree or building is still perceived as pointing up instead of sideways or down).

MacNeilage et al. (2010) compared visual and vestibular heading discrimination and found no effects of movement in head or world coordinates for visual heading in contrast to vestibular heading where thresholds were lower for horizontal translations when upright and also both visual and vestibular heading thresholds were consistently lower in the upright orientation compared to side-down for both horizontal and vertical plane movements. This suggests that vestibular heading is strongly dependent on movement in head coordinates and not world coordinates, meaning it is not influenced by or can compensate for the effects of gravity. However, vision appears to be even more specifically tuned to the upright orientation, suggesting that it may rely more on the longitudinal body axis as the preferred frame of reference. Claassen et al. (2016) had participants conduct a visual motion coherence task for vertical and horizontal motion while upright and when on their left side. In both body orientations, lower thresholds were found when the direction of the visual motion was congruent with the direction of gravity, but in different frames of reference - downwards motion while upright and rightward motion while left-side down. Although the second finding did not reach statistical significance, it suggests a potential difference in the frame of reference for the same task between the two body orientations. In the upright position, the visual motion may have been viewed as object motion in an earth-centered reference frame, whereas the left side down position was viewed as an egocentric reference frame (left side down produces otolith input analogous to rightward motion) (Claassen et al., 2016). In the current design, the yaw visual rotation was invariant to the gravity and idiotropic vector for the different body orientations used. Thus, the perceived timing of the visual motion may have been processed within an egocentric frame, relying primarily on the longitudinal axis signal and unable to account for the effect of gravity in an allocentric frame. This may potentially explain why the timing estimates were further from zero when the body was tilted from vertical compared to the upright orientation because gravity and the longitudinal axis are aligned in an upright posture but mismatched when the body was in all of the off-vertical orientations.

Why might the visual stimuli be processed in an egocentric frame and not combined into an allocentric representation to include the orientation of the body when vestibular and body signals are unreliable? Part of the reason may be from top-down influences and the context of the visual stimuli. In a study by Vidal et al. (2006), participants had to estimate the perceived angle of the bend of a curved virtual corridor presented using optic flow in the pitch and yaw axes while upright and lying right side down. In the upright position, forward pitch turns had significantly greater overestimation errors than backwards pitch and in the right side down position, estimation of yaw rightward turns were being overestimated more than left turns. This combination of rotation direction along with the respective body orientations represented the context equivalent to the direction of falling forward. This suggests that topdown influences from scene navigability and fear of falling with the direction of gravity may have led to the enhanced processing of visual motion resulting in the overestimation of the visually derived curve (Vidal et al., 2006). In a similar study using the same task, it was reported that the overestimation of downwards pitch turns were diminished when the task was performed while free floating in space compared to on earth (De Saedeleer et al., 2013). In support of the top-down influence hypothesis, this reduction in asymmetry could have been explained by the absence of risk of falling due to the weightlessness from free floating. In the current results, we did not find a significant difference for the direction of the visual yaw rotation for both right and left recumbent positions. This could be explained by the lack of orientation in the starfield environment that was used for the visual motion. Future work using more visually enriched content could be used to assess the effect of visual polarity cues.

Visual context has also been reported to influence the temporal perception of visual motion. Estimations of time-to-passage (TTP) are overestimated when accelerating in a vertical downwards direction (free fall) compared to acceleration vertical upwards and in an earth-horizontal direction (Indovina et al., 2013). Precision was also lower during constant acceleration in the vertical downward compared to acceleration suggesting that there is an expectation or anticipatory effect due to the presence of visual gravity (Indovina et al., 2013). Another top-down influence is a phenomenon called the visual reorientation illusion (VRI), which is when a person’s perception of upright is recalibrated based on the visual context (Oman, 2007). McManus et al. (2021) hypothesized that experiencing a VRI would lead to different perception of visual motion for prone and supine orientations due to reinterpretation of the gravity vector in an active visual distance estimation task. A VRI while supine would lead to gravity being ambiguous with a forward translation, therefore increasing visual gain or require less visual motion to reach a target and vice versa with prone orientation. However, this hypothesis was not supported by their findings that visual gain increased in both lying down orientations. Instead, they found that an increase in visual gain was correlated with the level of VRI which was largely dependent on whether the visual environment had orientation cues (a hallway vs. a starfield) (McManus and Harris, 2021). These results support a cognitive (top-down) hypothesis where the interpretation of the environment is more important with the level of VRI being correlated with the visual gain regardless of body orientation.

In addition to visual context, other characteristics of visual stimuli, such as field of view (FOV), disparity and scene layout, have also been reported to explain some of the differences observed in the bias of visual heading discrimination moderated through vection (de Winkel et al., 2018). Tanahashi et al. (2012) also found that increasing visual stimulus size increases vection strength induced by circular rotation around the pitch, yaw and roll axes. In the current study, visual stimuli was presented using a relatively new head mounted display (HMD) model with a limited FOV (Oculus Quest). It is possible that a larger display providing a more expansive FOV could generate more compelling and effective visual motion stimuli. The findings discussed so far have also been from a variety of apparatuses such as different models of HMDs, shrouded displays, as well as projector screens. Whether this interacts with how the visual context is perceived and the effectiveness of the visual motion can potentially explain some of the differences found across studies.

While these findings have demonstrated the varying contributions of vision, gravity and vestibular cues, as well as body orientation cues in the perception of upright align with previous reports (Jenkin et al., 2004). The question that remains is what are the mechanisms that govern the role of these factors in determining upright relative to the perception of motion? One potential mechanism that has been suggested that could explain these findings is the Bayesian optimal cue integration and sensory reweighting theory (Fetsch et al., 2009; Knill and Pouget, 2004). The idea behind the optimal integration and reweighting concept is that multiple sensory cues are combined based the relative reliability of the unisensory signal to determine the “weight” or contribution of that signal to the final representation. Miwa et al. (2019) reported that the longitudinal body axis dominates the perception of subjective uniform motion of a circle stimulus in the fronto-parallel plane when the direction of the visual motion matched the gravitational cue (upright position) without much consideration for visual polarity cues. Only when the body orientation cue was dissociated from gravity did visual polarity have an effect, supporting a Bayesian framework of shifting the representation of upright to rely more on the visual cues when the reliability of the gravity and body orientation cues were reduced due to conflict.

Interestingly, when the body orientation cue was dissociated from gravity and no visual cues were present, the perception still followed the longitudinal axis even when gravity cues (from vestibular input) were in conflict, suggesting a prior effect of relying on the longitudinal axis when no other cues of upright are available. This may explain our finding of why the perceived timing of visual motion was further from zero in the off-vertical orientations because no visual orientation cues were present, the timing estimate relied on the vestibular and longitudinal axis which were aligned in the upright orientation, but unreliable due to the mismatch when the body was tilted. Similar findings have been reported between the interaction of body and visual orientation cues, in which there is no influence of vision when upright and only when body orientation and gravity do not align, providing further support of the reweighting hypothesis (De Saedeleer et al., 2013; Hummel et al., 2016; McManus and Harris, 2021). Review of brain imaging studies on the reciprocal inhibitory interaction between visual and vestibular stimulation provides potential physiological evidence for the sensory reweighting mechanism when there is a sensory mismatch or incongruent sensory signals [see (Brandt et al., 2002) for a review]. Future research should include visual orientation cues, to determine whether the perceptual timing estimate will shift towards the visual signal when the vestibular and body cues are unreliable.

## Conclusion

In this study, we explored whether the presence of a mismatch in body orientation and vestibular cues due to gravity would influence the perceived onset of a visual yaw rotation. In all conditions we found that the onset of visual motion had to be presented before an auditory tone to be perceived as occurring simultaneously (127-159ms). We also found that visual motion had to be presented approximately 30ms earlier when an observer was in an off-vertical orientation compared to when they were upright for the stimuli to be perceived as occurring simultaneously. This suggests that the perception of upright signalled by the vestibular and body cues relative to gravity had a significant influence on the accuracy of responses in which there is a delay in the ability to detect visual motion when the body is tilted from upright. This additional delay may be attributed to the uncommon experience of motion when in a non upright posture or from the sensory conflict resulting from the dissociation between the vestibular and body cues relative to gravity, leading to a decrease in the reliability of these signals. Future work will need to include orientation cues in the visual stimuli and further control for the reliability of the gravity, body orientation and visual cues to further explore the interaction between the perception of upright and motion.

